# HumCFS: A database of fragile sites in human chromosomes

**DOI:** 10.1101/231233

**Authors:** Rajesh Kumar, Gandharva Nagpal, Vinod Kumar, Salman Sadullah Usmani, Piyush Agrawal, Gajendra P.S. Raghava

**Author notes:** Authors contributed equally to the work (Rajesh Kumar, Gandharva Nagpal). Corresponding authors: Gajendra P.S. Raghava, Center for Computational Biology, Indraprastha Institute of Information Technology, New Delhi 110020, India. Email of Authors: RK GN VK SSU PA.

## Abstract

Genomic instability is the hallmark of cancer and several other pathologies, such as mental retardation; preferentially occur at specific loci in genome known as chromosomal fragile sites. HumCFS (http://webs.iiitd.edu.in/raghava/humcfs/) is a manually curated database provides comprehensive information on 118 experimentally characterized fragile sites present in human chromosomes. HumCFS comprises of 19068 entries with wide range of information such as nucleotide sequence of fragile sites, their length, coordinates on the chromosome, cytoband, their inducers and possibility of fragile site occurrence i.e. either rare or common etc. Each fragile region gene is further annotated to disease database DisGenNET, to understand its disease association. Protein coding genes are identified by annotating each fragile site to UCSC genome browser (GRCh38/hg38). To know the extent of miRNA lying in fragile site region, miRNA from miRBase has been mapped. Comprehensively, HumCFS encompasses mapping of 5010 genes with 19068 transcripts, 1104 miRNA and 3737 disease-associated genes on fragile sites. In order to facilitate users, we integrate standard web-based tools for easy data retrieval and analysis.

## Introduction

The chromosomal fragile sites (CFS) are the specific chromosomal regions that exhibit an increased frequency of gaps and breaks when cells are exposed to DNA synthesis inhibitors^1^. These are the specific heritable loci, enriched with AT repeat sequences, and are evolutionarily conserved^2^. The fragile site can be classified as rare and common according to their frequency of expression in the population^3^. Rare fragile sites are present in a small fraction of the population (<5%), while the common fragile sites are basic components of chromosomes, as these sites are present in approximately all individuals^4^. Fragile sites are said to be expressed when they exhibit cytogenetic abnormality i.e. appears as gaps or breaks in metaphase chromosome and can extend up to millions of base pair (Mbp) in length^5^. Rare fragile sites are specifically induced by BrdU (Bromodeoxyuridine) and folic acid thus could be sub-categorized as BrdU (Bromodeoxyuridine) sensitive and folate-sensitive. In case of common fragile sites, Aphidicolin, 5-Azacytidine, and Distamycin-A are most common inducers^3^.

Over the past few years, fragile sites have been realized to be an important aspect of cancer biology, as most of the cancer-related genes occur in the CFS^6^. The instability of fragile sites often results in aberrant expression of oncogenes and tumor-suppressing genes, a step towards initiation of cancer progression^7^. It has also been shown by in vitro studies that translocation, deletion, intra-chromosomal gene arrangement and sister chromatid exchange of cancer-specific genomic regions occur as a consequence of cell treatment with fragile site inducers^8^and common fragile sites even also co-localize with breakpoints and deletions specific to various tumors as shown in in vitro cancer cell line studies^9,10^. Tumor suppressor gene e.g. WWOX located within the FRA16D fragile site is often aberrantly methylated and can be correlated with the development of various tumors such as ovary, prostate, and breast cancer^11,12^. The aberrant expression of WWOX gene is because of hyper methylation at CpG Island that may have a correlation with chromosomal common fragile sites being rich in CGG/CCG trinucleotide^6^. Micro-RNA (miRNA), which is essential for cell survival, cell differentiation, metabolism and cell death has also been reported in fragile sites. e.g. FRA4D contains miR-218-1 and FRA5G contain miR-218-2^6^. The deregulated expression of miRNA due to chromosomal rearrangement has been associated with cancer-specific events and tumor development^13,14^. e.g. the differential expression of miR218 due to chromosomal rearrangement is associated with epithelial cancer development^7^.

In the past numerous resources has been developed to maintain wide range of information related to instabilities in genome and chromosomes that includes i) TICdb: contain information on chromosomal translocation breakpoint in human tumors^15^, ii) HYBRIDdb: maintain information of hybrid genes in humans^16^, iii) dbCRID: compile information on chromosomal rearrangement in diseases^17^and iv) chimerDB3.0: maintain fusion genes^18^. Since from the discovery of CFS, several lines of evidence suggest their involvement in human disease progression, as they are preferred sites for exogenous DNA insertion, chromosomal translocation, re-arrangement, and breakpoint. But there remains a large gap in our knowledge about human CFS and its association with various diseases. One possible reason could be the unequal distribution of divergent studies in literature, which imposes a problem in the comparative study of these versatile human genomic regions. Best of our knowledge there is no database in the literature that maintains information on human chromosomal fragile sites despite the fact that fragile sites are core genomic regions responsible for instability and diseases. Keeping this view in mind and to complement other related existing resources we developed a database related to human chromosomal fragile sites.

## Results

### HumCFS database statistics and significant findings

HumCFS is a unique repository of human CFS and associated data related to gene-disease relationship and miRNA with them. The database includes 118 experimentally verified molecularly cloned human chromosomal fragile sites, which are manually curated from literature for the entire somatic chromosome except for the sex chromosomes X and Y. By calculating the nucleotide length of each fragile site compared to the non-fragile site, we found that fragile sites constitute 19% of the total human genome length. The database contains entries of 5010 protein-coding genes, which coincide with the fragile sites of human genome compared to 14248 manually curated protein-coding genes available in ensemble genome file of Havana protein-coding genes. This suggests that 35.16% of protein-coding genes overlap with the fragile regions of the human genome. This indicates that fragile sites are rich in an essential genome. The number of fragile sites, genes, and miRNAs corresponding to each chromosome is shown in Figure 1. When we annotate the fragile sites with respect to miRNA, we found that 1104 miRNAs are encoded from region lying in the fragile sites, which corresponds to approximately 42.77% of total miRNA present in miRBase. This reaffirms an important observation that overall distribution of regulatory part of the genome is much higher in fragile sites. DisGenNET is one of the largest publicly available collections of genes and variants associated to human disease^19^. Out of 5010 protein-coding genes found to be present in human CFS in the current study, we were able to map 3737 (74.5%) genes to DisGenNET database using HGNC symbols, indicating their association with human maladies. A higher number of genes are found to associate with neoplasm, nervous system disease, pathological conditions and mental ailments. The distribution of these genes among various diseases classes recognized by DisGenNET is shown in Figure 2. We observe that chromosome 1, 2, 10, 11, 12, 19 shows a higher percentage of protein-coding genes in fragile sites while 9, 13, 14, 15, 16, and 17 shows less percentage of the protein-coding gene. We also observed that chromosome number 2 and 19 have the highest number of protein-coding genes miRNAs in fragile regions as compared to all another chromosome. We also observed that Aphidicolin is the most potent inducer of chromosomal fragility while DiatamycinA is the least potent inducer.

**Figure 1:**
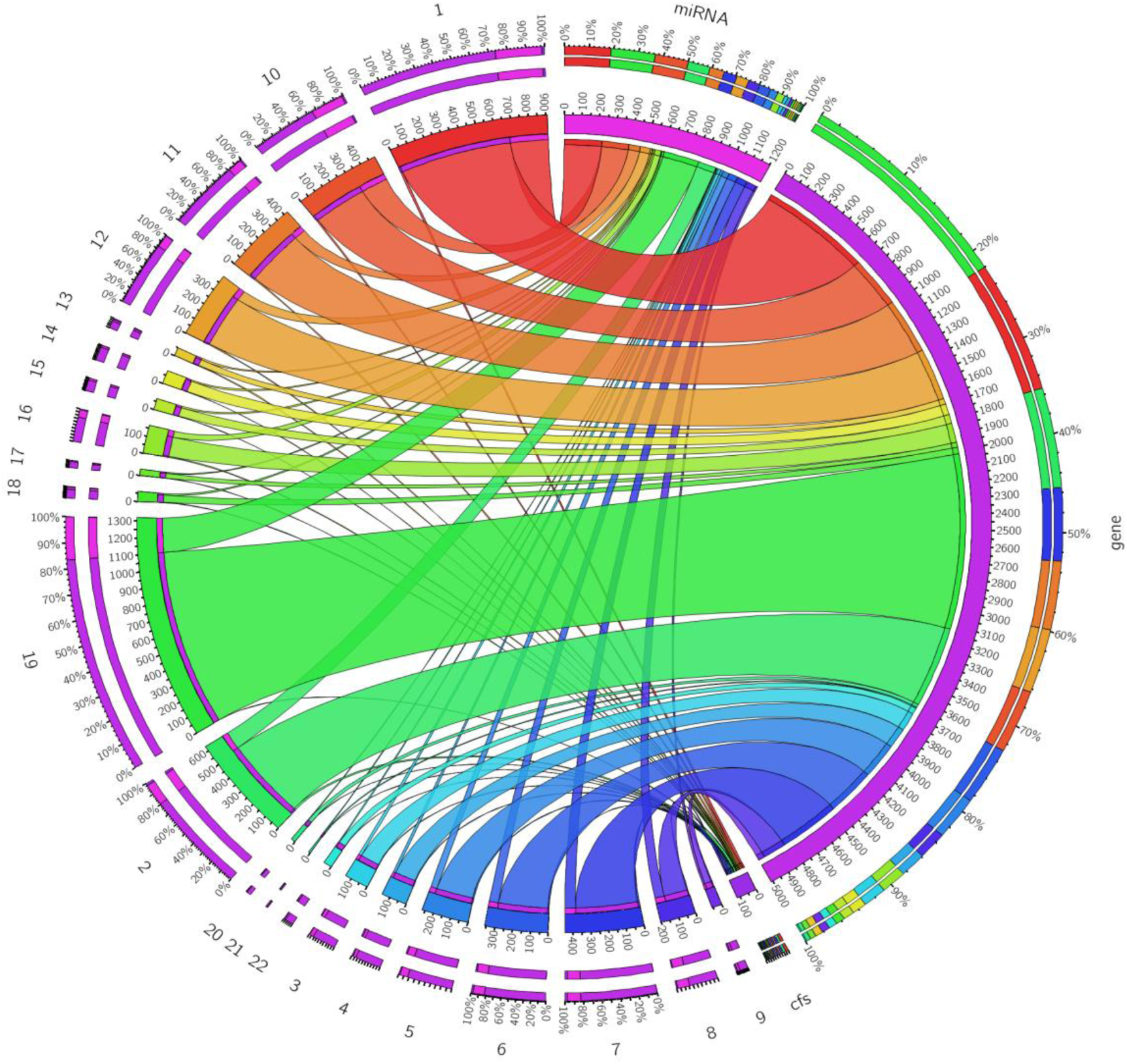
Circos diagram explaining the number of fragile sites, genes, and miRNA in each chromosome. ( letter 1-22 denotes chromosome number, cfs denote chromosomal fragile site.

**Figure 2:**
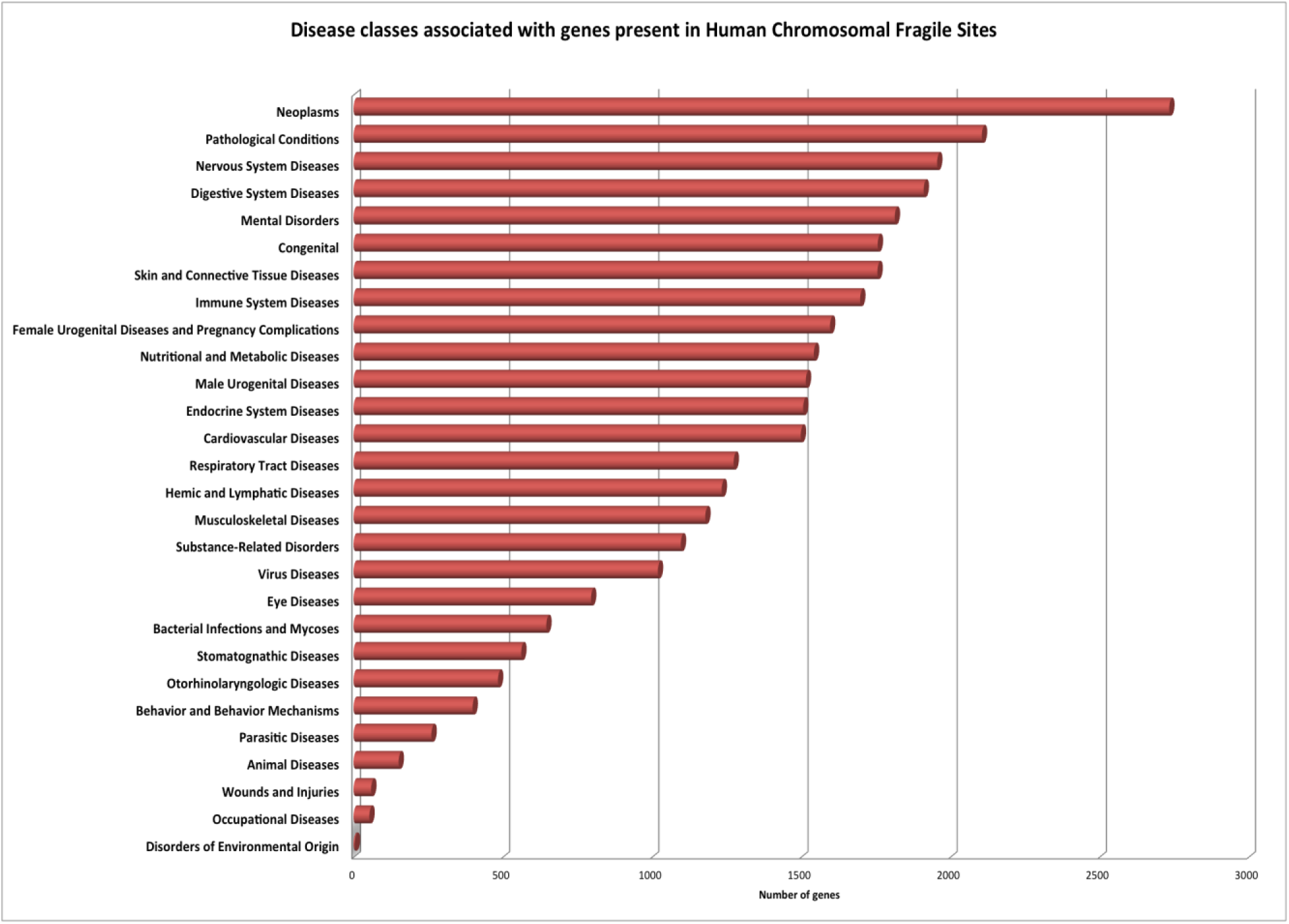
Distribution of genes among various diseases classes recognized by DisGenNET.

## Discussion

Genomic instability in the form of chromosomal rearrangement and mutation is the characteristics of almost all type of cancer^20–22^. In genome, two types of factors play an important role in genome instability-one that acts in trans and suppresses genome instability, which includes damage repair mechanism and cell cycle checkpoint inhibitors and second is the chromosomal hotspots for the genomic instability known as fragile sites which are AT-rich sequences, evolutionarily conserved and are highly transcribed^23,24^. But the knowledge about human CFS is scattered in literature, posing challenges for studying these important regions of the human genome. Keeping in view the necessity of unified platform, an attempt has been made in the present study to develop a knowledgebase for exploring the human chromosomal fragile sites. In HumCFS, all relevant information regarding chromosomal fragile sites has been compiled in a systematic manner, which will help the researchers to look into a variety of aspects of this important region of the genome. In the present study information regarding chromosomal fragile site has been manually curated from research articles and annotated using Ensembl gene files, miRBase and DisGenNET database which provide comprehensive information for each of the entry made in the database. We observed chromosome 2 and 19 have the highest number of protein-coding genes and miRNA which is consistent with the previous study^6^ and chromosome 13, 14 and 15 shows the lowest number of miRNA and protein-coding genes. This shows that distribution of genetic element and regulatory element in the genome is not even, it depends upon the chromosome. We also observed that most of the fragile sites in human are induced by Aphidicolin, an important finding that never been in the literature so far while Distamycin –A is the least potent inducer. Briefly, the user can take benefit from HumCFS in following ways (i) the user can browse a fragile site with well annotation by one click, this will save time (ii) extract the information about moonlight disease (iii) visualize genome by genome browser. In summary, HumCFS is a useful resource that will expedite the human chromosomal fragile site research.

The current study is an initiative that could pave way for prenatal prediction of possible health challenges that could be incurred by the individual due to the chromosomal breakage events. In-depth understanding of the relationship of fragile sites with diseases is a prerequisite for the determination of therapeutic strategies based on the genomic profile of an individual. Thus, in future, detection of chromosomal breakage along with the genomic site of the breakage event could become a part of the genomic profiling of patients that could help in making the choice of disease management. Although the present study is a comprehensive resource, anticipated to provide an impetus to the fragile site-disease association research and application; diligent efforts are required to apply this knowledge in the prognosis of diseases. This would be possible with the rigorous disease-specific investigation of associated chromosomal breakage events. In literature most of the study links CFS to cancer but we think that there is need to shift this paradigm to other disease related study also, as genes related to cardiovascular, metabolic, mental, musculoskeletal, respiratory, nervous system etc. are also found in human CFS.

## Material and Methods

### Data collection and compilation

In order to collect the latest information about fragile sites an extensive search was performed in PubMed by using the combination of keyword string such as ‘fragile site’, ‘chromosomal fragile region’, ‘Human chromosomal fragile site’, ‘Fragile region of human’, ‘Chromosomal fragile site’. This search results into 4714 PubMed abstract, then each abstract is manually curated for relevant information regarding human CFS, which includes fragile site name, its coordinate, cytoband, inducer, frequency and its type. Relevant information, which is included in the database, comes from 65 PubMed articles, which contains information about molecularly cloned experimentally verified human CFS. Region corresponds to chromosomal coordinate as given in PubMed articles is considered as fragile site and rest of the region of the chromosome is considered as non-fragile region of the genome. Further to provide the complete and comprehensive information regarding the human chromosomal fragile site we have done three level of curation, Firstly, the information regarding protein-coding genes lies within human CFS was annotated using UCSC genome browser (GRCh38/hg38)(http://hgdownload.soe.ucsc.edu/goldenPath/hg38/chromosomes/) by using in-house PYTHON scripts. Secondly, miRNA associated with each gene present in human CFS was annotated from miRBase^25^. Thirdly, genes present in human CFS, which are associated with the human ailment, are annotated by using DisGenNET database. Finally, 5010 protein-coding genes systematically and comprehensively compiled in HumCFS.

### Database architecture and web interface

HumCFS database has been built using a standard platform based on the Linux-Apache-MySQL-PHP (LAMP). Red Hat Linux (version 6.2) as the operating system, MySQL (version 14.12) for managing the data and Apache (version 2.2.17) as the HTTP server was used for developing this database. HTML5, PHP, JAVA scripts have been used for developing the mobile and tablet compatible front ends and MySQL for developing the back end. For the entire database interface and the common gateway, networks PHP and PERL coding have been used. The complete architecture of HumCFS database is given in Figure 3.

**Figure 3:**
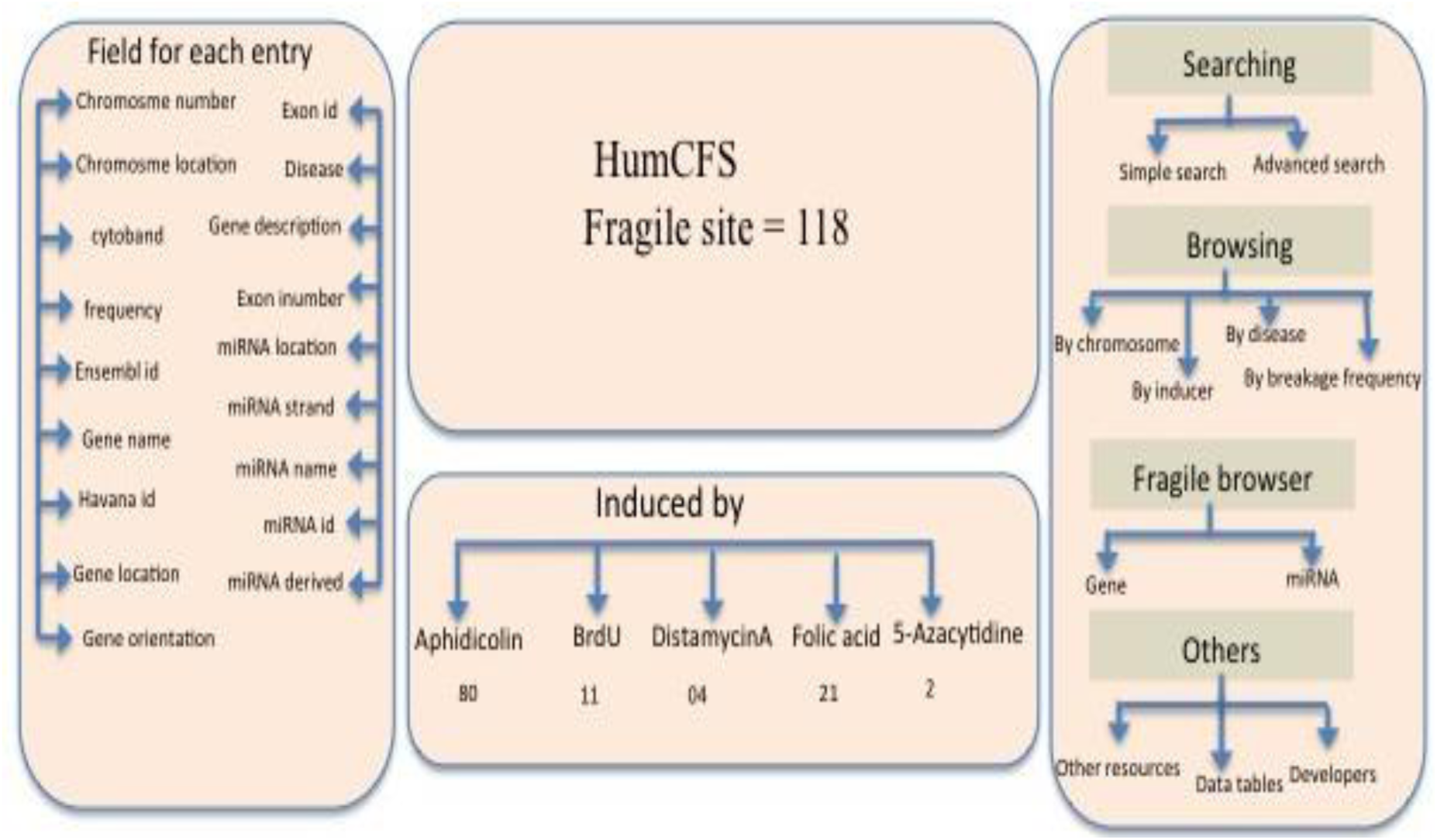
A Schematic representation of HumCFS database architecture.

### Data organization

Primary information regarding human CFS was manually curated from PubMed. Each entry includes following information: name of the fragile site, chromosomal location, cytoband, type, frequency, nucleotide sequence and chromosomal number. Apart from the primary information, secondary information related to gene name, gene ensemble ID, gene location, gene orientation, nucleotide sequence, transcript number, and ID was annotated from UCSC genome browser and maintained in a separate entry which provides a smooth and flawless searching and data retrieval while using the database. miRNA information like miRNA name, miRNA ID, miRNA location, strand information, related to each gene is annotated using miRBase and maintained in database. Lastly, eachprotein-coding gene associated with human disease is annotated by using DisGenNET database; hence provide a complete and well-annotated picture of the HumCFS database.

### Implementation of tools

#### Search tools

The search tools which allows users to perform their query by performing a search in any field of the database like fragile site name, cytoband, chromosomal coordinate, type of inducers, miRNA associated with the gene. This module allows a user to search selected or all field of the database for selected search record.

#### Browse tools

HumCFS is equipped with browsing facility that allows accessing data on major field includes (1) By chromosome (2) By fragile site inducer (3) By frequency of breakage (4) Moonlight disease search. Data according to chromosome wise can be retrieved and can be compared, which provides information about which chromosome contains the highest number of gene in fragile sites, this allows users to refine their work. Browsing by inducer provides the additional facility to user look for human CFS induced by the particular chemical. Breakage frequency module allows the user to categories fragile site in common and rare ones. Many genes present in HumCFS linked to more than one functional categories of a disease known as moonlight property. Therefore to aid in search of this kind of moonlight gene, we have provided ‘Moonlight disease’ function in our database. The user may search for a gene, which in addition to link with cancer is also linked with cardiovascular and tuberculosis disease etc. So this search criterion allows the user to perform a search for multi-functional gene pairs.

#### Sequence alignment

In order to perform sequence similarity based search, BLASTN^26^is integrated into the HumCFS database. The user can submit their nucleotide sequence in FASTA format up to 10-1000 lengths. The server performs BLASTN search for the user’s query sequence against the nucleotide sequences of all the fragile sites present in the database.

#### Genome browser

We also integrated an interactive ‘Genome browser’ who is powered by JBrowse^27^a JavaScript and HTML5.0 based browser to develop descriptive section using JSON (JavaScript Object Notation data format) which allows fast, smooth, scrolling of fragile site genomic data with unparalleled speed. By clicking on gene name or miRNA name all the information regarding that particular entity including sequence; location, ID, sequence etc. can be retrieved from the JBrowse Figure 4.

**Figure 4:**
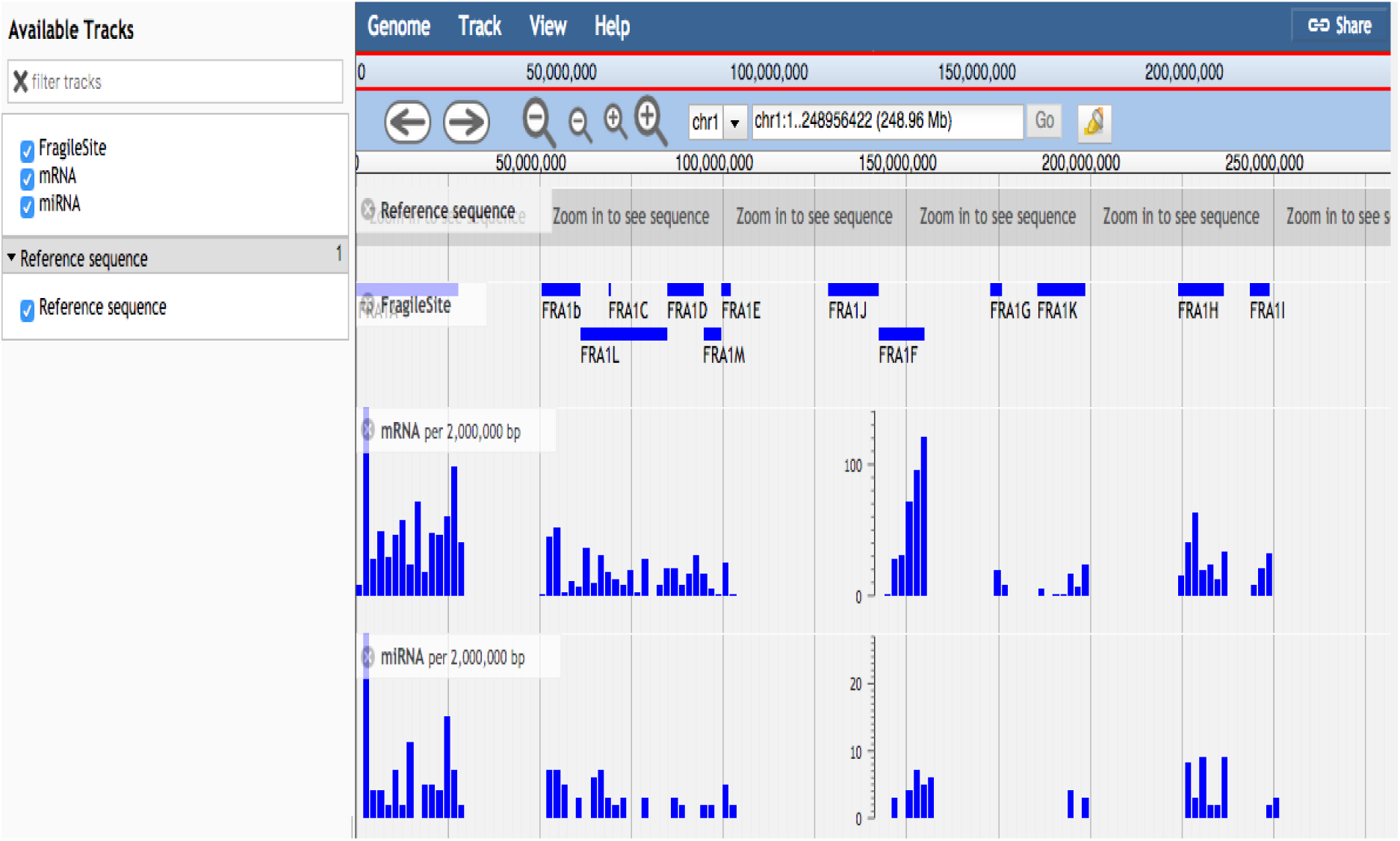
Figure describing data visualization using genome browser.

## Availability

http://webs.iiitd.edu.in/raghava/humcfs/

## Acknowledgements

Authors are thankful to funding agencies, Council of Scientific and Industrial Research (CSIR), University Grant Commission (UGC), Department of Science and technology (DST) and Department of Biotechnology (DBT), Govt. of India for financial support and fellowships.

## Author contributions

RK and GN manually and programmatically collected and curated all the data. RK, GN analyzed the data. RK, VK and SSU developed the web interface. SSU and PA helps in file processing. RK, GN, SSU and GPSR prepared the manuscript. GPSR conceived the idea, planned and coordinated the entire project.

